# Lessons in effector and NLR biology of plant-microbe systems

**DOI:** 10.1101/171223

**Authors:** Aleksandra Białas, Erin K. Zess, Juan Carlos De la Concepcion, Marina Franceschetti, Helen G. Pennington, Kentaro Yoshida, Jessica L. Upson, Emilie Chanclud, Chih-Hang Wu, Thorsten Langner, Abbas Maqbool, Freya A. Varden, Lida Derevnina, Khaoula Belhaj, Koki Fujisaki, Hiromasa Saitoh, Ryohei Terauchi, Mark J. Banfield, Sophien Kamoun

## Abstract

A diversity of plant-associated organisms secrete effectors—proteins and metabolites that modulate plant physiology to favor host infection and colonization. However, effectors can also activate plant immune receptors, notably nucleotide-binding domain and leucine-rich repeat-containing (NLR) proteins, enabling plants to fight off invading organisms. This interplay between effectors, their host targets, and the matching immune receptors is shaped by intricate molecular mechanisms and exceptionally dynamic coevolution. In this article, we focus on three effectors, AVR-Pik, AVR-Pia, and AVR-Pii, from the rice blast fungus *Magnaporthe oryzae* (syn. *Pyricularia oryzae*), and their corresponding rice NLR immune receptors, Pik, Pia, and Pii, to highlight general concepts of plant-microbe interactions. We draw 12 lessons in effector and NLR biology that have emerged from studying these three little effectors and are broadly applicable to other plant-microbe systems.

## INTRODUCTION

In the last ~10 years, the field of effector biology has played a pivotal role in the study of plant-microbe interactions. The current paradigm is that unravelling how plant pathogen effectors function is critical for a mechanistic understanding of pathogenesis. This concept applies to pathogens and pests as diverse as bacteria, fungi, oomycetes, nematodes and insects. In addition, plant symbionts are also known to secrete effectors for optimal interactions with their hosts. Much progress has been made in describing the diversity of effector genes from pathogen genomes, understanding how effectors evolve and function, and applying this knowledge to real-world problems to improve agriculture.

The field of effector biology is also intimately linked to the study of plant immunity. Plants are generally effective at fighting off invading pathogens. They have evolved immune receptors that detect pathogen effectors to activate effective defense responses (Cesari et al. 2014a; Dodds and Rathjen 2010; Jones and Dangl 2006; Takken and Goverse 2012; Van Der Hoorn and Kamoun 2008; Win et al. 2012a). Among these receptors are the nucleotide-binding, leucine-rich repeat proteins (NLR or NB-LRR), an ancient class of multi-domain receptors also known to confer innate immunity in animals (Duxbury et al. 2016; Maekawa et al. 2011). More recently, a subset of NLR proteins were discovered to carry extraneous domains that, in some cases, have evolved by duplication of an effector target, followed by fusion into the NLR. This finding further cements the merger of two major fields of plant-microbe interactions—effector biology and NLR biology—with important fundamental and practical implications.

Plant diseases are a recurring threat to food production and a major constraint for achieving food security (Bebber et al. 2014; Bebber and Gurr 2015; Fisher et al. 2012; Pennisi 2010). A prime case is blast, a disease caused by the ascomycete fungus *Magnaporthe oryzae* (syn. *Pyricularia oryzae*), which is best known as the most destructive disease of rice. In addition to rice, *M. oryzae* can infect other cereal crops such as wheat, barley, oat and millet destroying food supply that could feed hundreds of millions of people (Fisher et al. 2012; Liu et al. 2014; Pennisi 2010). Increased global trade, climate change, and the propensity of this pathogen to occasionally jump from one grass host to another, have resulted in increased incidence of blast diseases.

The completion of the genome sequence of the rice blast fungus *M. oryzae*, about 12 years ago, ushered in an exciting era of genomics research for fungal and oomycete plant pathogens (Dean et al. 2005; 2012; Dong et al. 2015). Through the following decade, plant pathogen genome sequences became a unique resource for basic and applied research, enabling many creative approaches to investigate both pathogen and plant biology (for examples, see Pais et al. 2013). In particular, a number of *M. oryzae* effectors have been discovered and validated, primarily through their avirulence (AVR) activity—the capacity to activate immunity in particular host genotypes. Among these are the three effectors AVR-Pik, AVR-Pia, and AVR-Pii corresponding to three rice NLR pairs, Pik, Pia, and Pii (Yoshida et al. 2009) (Fig. 1, Table 1). The objective of this article is to highlight lessons drawn from knowledge of the structure, function and evolution of these three little effectors of the rice blast fungus and their matching rice NLRs. In total, we list 12 lessons in effector and NLR biology that are broadly applicable to other plant-microbe systems and could serve as a benchmark for future conceptual developments in effector and plant immunity research.

**Figure 1.**
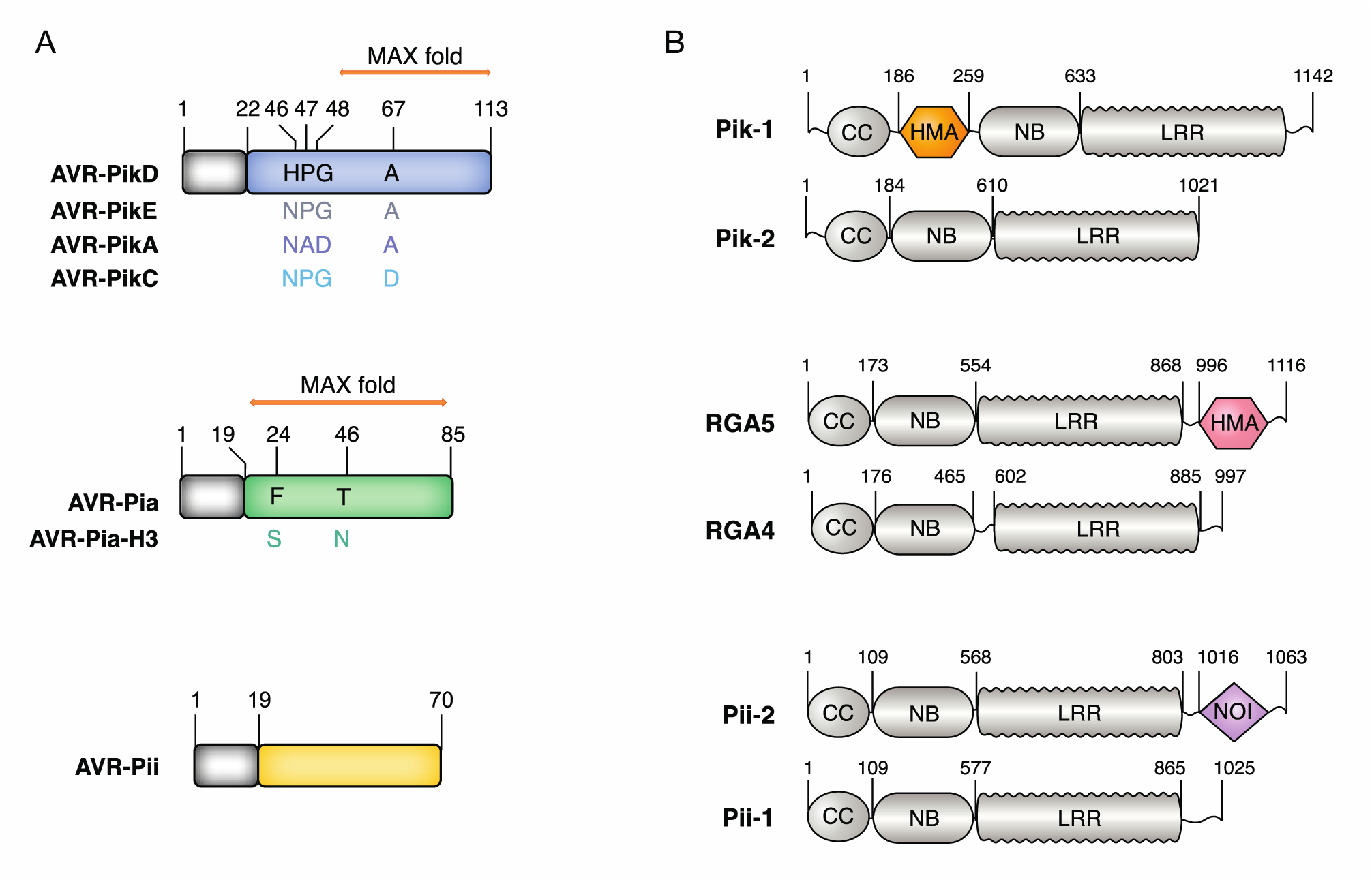
Three little effectors and corresponding plant NLRs. (A) The alleles of the *Magnaporthe oryzae* effectors AVR-Pik, AVR-Pia, and AVR-Pii showing the position of polymorphic residues. The effector domain is shown in color, while the signal peptide is in grey. The conserved MAX fold in AVR-Pik and AVR-Pia is displayed as an orange arrow. (B) Rice Pik, Pia (RGA5/ RGA4), and Pii NLRs highlighting the position of integrated HMA (hexagon) and AvrRpt cleavage/NOI (diamond) domains in the classical plant NLR architecture (CC, coiled coil; NB, Nucleotide-binding; LRR, leucine rich repeat domain).

**Table 1.** An overview of the rice blast effectors AVR-Pik, AVR-Pia and AVR-Pii and their matching NLRs.

| Effector | Accession number | Structure | Matching NLR pair | Accession number | Integrated domain | Mechanism |
| --- | --- | --- | --- | --- | --- | --- |
| AVR-Pik | AB498875 | X-ray crystallography (Maqbool et al. 2015) | Pik-1 | HM035360 | HMA* (Maqbool et al. 2015) | Cooperation |
|  |  |  | Pik-2 | HM035360 | No |  |
| AVR-Pia | AB498873 | NMR (de Guillen et al. 2016) | Pia-1 (RGA4) | AB604622 | No | Negative regulation |
|  |  |  | Pia-2 (RGA5) | KC777364 | HMA* (Cesari et al. 2013) |  |
| AVR-Pii | AB498874 | Not available | Pii-1 (Pi5-1) | EU869185 | No | Not known |
|  |  |  | Pii2 (Pi5-2) | EU869186 | AvrRpt cleavage/NOI# (Sarris et al. 2016) |  |

\*Heavy Metal Associated domain (HMA)
#Not validated experimentally

### Lesson 1. The power of diversity: beyond reference genomes

Comparative genomics is a powerful tool that can be used to clone pathogen effector genes. Using this approach, Yoshida et al. (2009) identified three novel *M. oryzae* effector genes with avirulence (AVR) activity (*AVR*-*Pik*, *AVR*-*Pia* and *AVR*-*Pii*) matching three previously described rice resistance genes (*Pik*, *Pia*, *Pii*) (Yoshida et al., 2009). Yoshida et al. (2009) first used genetic association, but failed to identify the AVR effectors in the *M. oryzae* reference genome, 70-15 (Dean et al., 2005). An assessment of the 70-15 strain revealed that it has weak virulence, causing intermediate responses on the rice cultivars screened. In fact, 70-15 is a hybrid laboratory strain derived from a cross between two blast fungus isolates; 104-3 isolated from rice (*Oryza* spp.) and AR4 isolated from weeping lovegrass (*Eragrostis curvula*), with subsequent backcrosses to rice infecting isolate Guy11 (Chiapello et al. 2015). It is now established that 70-15 carries a hybrid genome, given that the *Eragrostis* parent is not classified as *M. oryzae per se* (Chiapello et al. 2015). The 70-15 genome may, therefore, lack some of the AVR effectors detected by rice resistance genes and the ambiguous reactions observed on 70-15 infected rice cultivars hampered further genetic analyses (Yoshida et al. 2009). To address this issue, Yoshida et al. (2009) sequenced the genome of Ina168, a *M. oryzae* isolate from Japan, expected to carry nine AVR effector genes based on response to classic rice blast resistance genes. Association genetics of Ina168 with 21 additional strains, mainly from Japan, is what ultimately led to the identification of the three AVR effectors discussed in this article.

Similar to many filamentous plant pathogens, *M. oryzae* has a dynamic genome and exhibits a high degree of genetic diversity (Talbot et al. 2003; Raffaele and Kamoun, 2012). The Yoshida et al. (2009) paper highlights the importance of looking beyond so-called “reference genomes” to consider the wider genetic diversity of a given pathogen species. The capacity to generate robust phenotypic data is also a critical component of genetic analysis and an essential complement to the genomic data. Moreover, the strains selected for comparative genome analyses should be amenable to phenotyping and must display distinct phenotypes, e.g. virulence. As the pathogenomics community expands beyond reference genomes and simple comparative studies, it becomes critical to establish robust phenotyping platforms and to select strains that can be processed through these pipelines.

### Lesson 2. Missing in action: unstable genomes drive pathogen evolution

Filamentous pathogen genomes are often organized in a so-called “two-speed genome” architectures in which effector genes occupy dynamic, repeat rich compartments that serve as cradles for adaptive evolution (Croll and McDonald 2012; Dong et al. 2015; Faino et al. 2016; Haas et al. 2009; Raffaele and Kamoun 2012; Yoshida et al. 2016). In *M. oryzae*, effector genes are often located in telomeres and near repetitive sequences and transposable elements, all of which are thought to be hot-spots for deletion, translocation, and duplication (Dong et al. 2015; Yoshida et al. 2016). Indeed, loss of AVR effector genes is a predominant mechanism for *M. oryzae* to evade rice immunity, with *AVR*-*Pik*, *AVR*-*Pia*, and *AVR*-*Pii* all exhibiting notable levels of presence/absence polymorphisms in *M. oryzae* field isolates (Huang et al. 2014; Yoshida et al. 2009). All three effector genes are linked to genetic elements indicative of genomic instability: *AVR*-*Pik* is flanked by repetitive sequences, *AVR*-*Pia* is linked to a transposable element, and *AVR*-*Pii* is located in a telomeric region (Fig. 2) (Huang et al. 2014; Yoshida et al. 2009). Notably, *AVR*-*Pia* has translocated to different *M. oryzae* chromosomes, suggesting that its associated locus is prone to chromosome rearrangements (Sone et al. 2012; Yasuda et al. 2006). *AVR*-*Pia* may hitchhike with a neighboring retrotransposon to move across the genome and laterally between strains as has been proposed for another *M. oryzae* effector, *AVR*-*Pita* (Chuma et al. 2011). Such effectors have been coined mobile effectors by Chuma et al. (2011) and may persist within an asexual pathogen population despite recurrent adaptive deletions (Chuma et al. 2011; Raffaele and Kamoun 2012).

**Figure 2.**
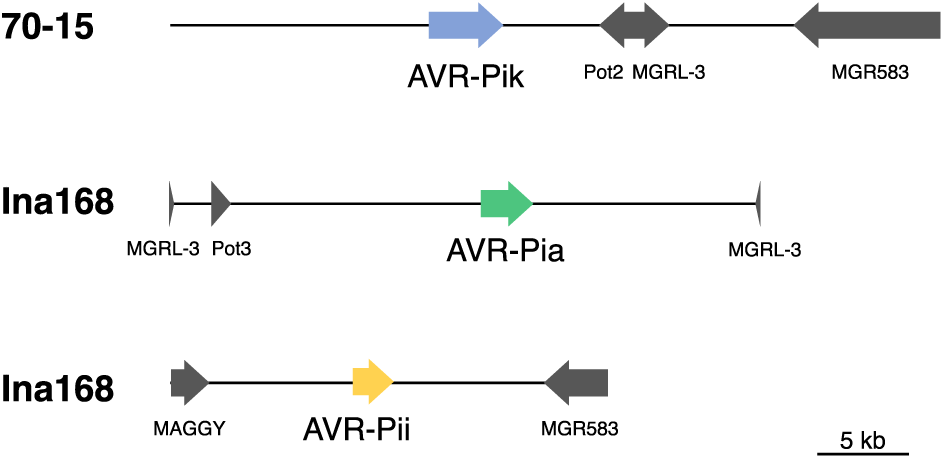
Linkage of *AVR*-*Pik*, *AVR*-*Pia* and *AVR*-*Pii* to transposable elements. The genomic regions of *AVR*-*Pik* (blue) in 70-15, *AVR*-*Pia* (green) and *AVR*-*Pii* (yellow) in Ina168 are shown, with arrowheads representing the direction of the coding strand. These three effectors are linked to various transposable elements (grey).

Deletions of AVR effectors in filamentous plant pathogens in response to host immunity provide unambiguous examples of rapid evolutionary adaptations that arise over short evolutionary timescales (Inoue et al. 2017). Beyond the instances described here, there are dozens of cases of AVR effector gene deletions in fungal and oomycete plant pathogens (Dong et al. 2015; Raffaele et al. 2010; Rouxel and Balesdent 2017; Sharma et al. 2014). In many cases, the effector genes occupy unstable, repeat-rich regions of the genomes that exhibit higher rates of structural variation. The capacity of the “two-speed genomes” of filamentous plant pathogen to accelerate evolution—and thus the pathogen’s capacity to adapt to the host immune system—is now well established as a fundamental concept in plant pathogenomics. However, the precise mechanisms by which genome architecture favors the emergence of new pathogen races requires further investigation (Croll and McDonald 2012; Dong et al. 2015; Faino et al. 2017; Raffaele and Kamoun, 2012).

### Lesson 3. The X Games: effector genes with extreme signatures of selection

In addition to presence/absence polymorphisms, many filamentous pathogen effectors show high levels of non-synonymous (amino acid replacement) sequence substitutions relative to synonymous changes (Allen et al. 2004; Dodds et al. 2004; Huang et al. 2014; Liu et al. 2005; Raffaele et al. 2010; Yoshida et al. 2009). Such patterns of sequence polymorphisms are a hallmark of positive selection and are thought to reflect the coevolutionary arms race of these effectors with host components. In some cases, the signatures of positive selection observed in effector genes are extreme, with only a few synonymous or no synonymous polymorphisms.

AVR-Pik is a striking example of an effector gene with an extreme signature of positive selection. Allelic variants of *AVR*-*Pik* exhibit four amino acid replacements, but not a single synonymous change (Huang et al. 2014; Yoshida et al. 2009) (Fig. 1A). Remarkably, all non-synonymous *AVR*-*Pik* polymorphisms are presumably adaptive, since they map to regions in the protein structure that interface with the Pik-1 immune receptor (Maqbool et al. 2015). In some cases, AVR-PikD polymorphisms have been shown to alter the binding affinity between the effector and the Pik-1 NLR protein, demonstrating the adaptive nature of these polymorphisms (Maqbool et al. 2015).

The excess of non-synonymous substitutions observed in effectors can make the computational identification of genes underlying adaptive evolution challenging, because similar patterns would arise from relaxed selection or in populations with small effective size (Terauchi and Yoshida 2010; Yoshida et al. 2016). This challenge is evident in *M. oryzae*, as about one third of candidate effector genes show outlier ratios of non-synonymous to synonymous substitutions (Yoshida et al. 2016). However, as the case of AVR-Pik clearly illustrates, many examples of extreme patterns of nonsynonymous substitutions in effector genes are probably adaptive, resulting from the stringent selection pressure conferred by disease resistant hosts (Win et al. 2007; Yoshida et al. 2016).

### Lesson 4. Appearances can be deceiving: sequence unrelated effectors share similar folds

Filamentous plant pathogen effectors are typically small secreted proteins that exhibit rapid levels of sequence diversification. One of the hallmarks of effector proteins is that they tend to have orphan sequences with little similarity to each other, or to known proteins. Interestingly, the threedimensional structures of effector proteins revealed unexpected similarities between unrelated proteins. For example, three-dimensional structures of the *M. oryzae* effectors AVR-Pia, AVR1-CO39, AVR-PikD, and AvrPiz-t contain the same core fold, despite sharing no apparent sequence similarity (de Guillen et al. 2015; Maqbool et al. 2015; Ose et al. 2015; Zhang et al. 2013) (Fig. 3).

**Figure 3.**
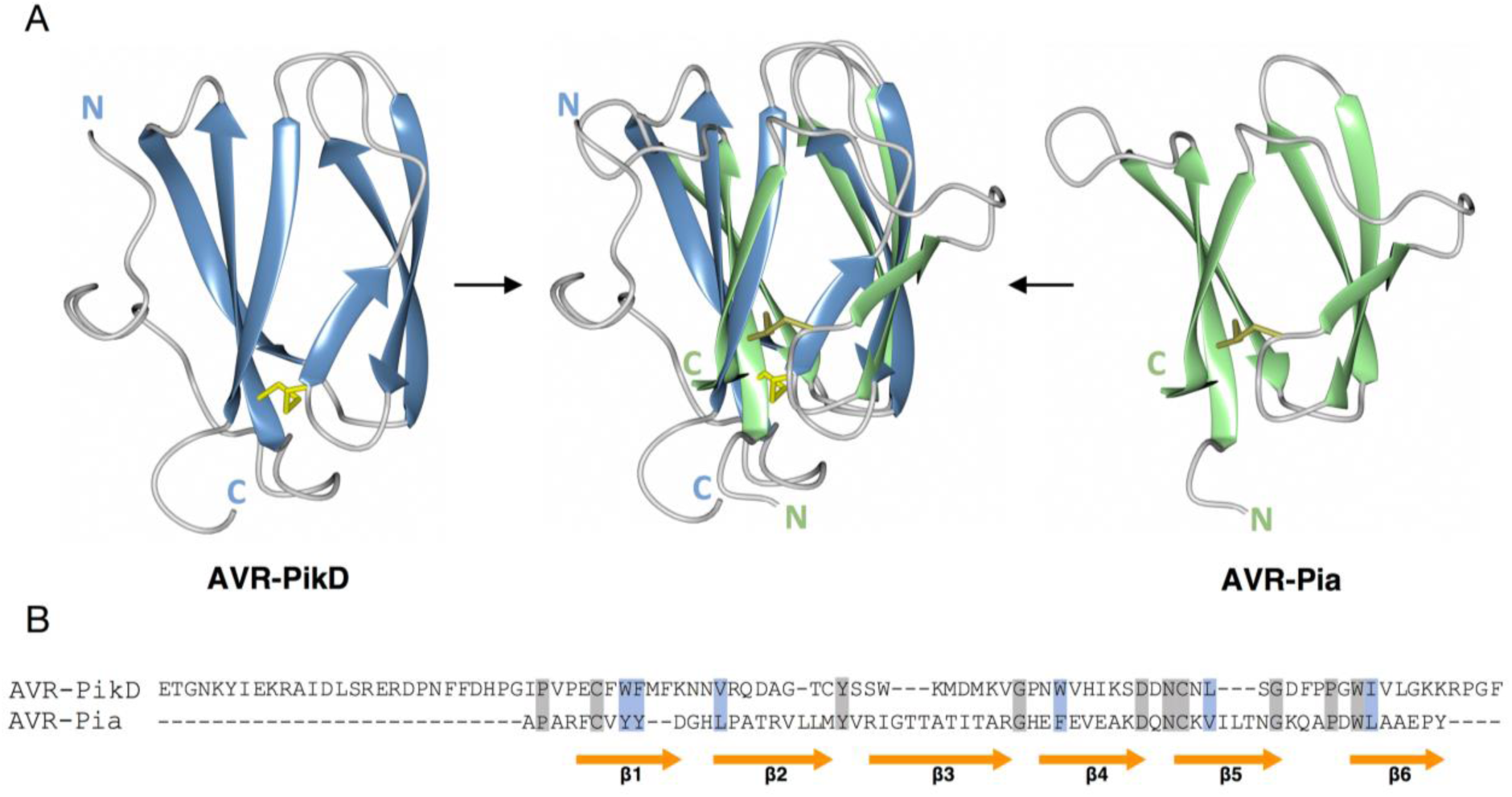
Sequence unrelated AVR-PikD and AVR-Pia share the same structural fold. (A) Schematic representation of AVR-PikD (left), AVR-Pia (right) and their superposition (centre), showing the conserved six strand β-sandwich MAX fold as a cartoon. Cysteines forming disulphide bonds are illustrated by yellow sticks, while flexible loops are in grey. Amino and carboxyl termini are labelled N and C, respectively. (B) Amino acid sequence alignment of AVR-PikD and AVR-Pia, highlighting the lack of conservation between the two effectors. Conserved amino acids are shaded in grey, amino acids with similar properties in blue. The structural conserved β-strands (β1 to β6) are depicted as arrows (orange).

Moreover, the host-selective toxin ToxB from *Pyrenophora tritici*-*repentis*, despite being derived from a phylogenetically distant pathogen, also contains the same fold—a six-stranded β-sandwich, comprised of two anti-parallel β-sheets (Nyarko et al. 2014). de Guillen et al. (2015) designated effectors containing this conserved β-sandwich core as “MAX effectors” (for *<u>M</u>. oryzae* <u>A</u>VRs and To<u>x</u>B) and identified additional family members in various ascomycete fungi. MAX effectors have particularly expanded in *M. oryzae* (5-10% of total effectors) and in the related species *Magnaporthe grisea* (de Guillen et al. 2015). The distribution and diversity patterns of the MAX effector family indicate that they expanded through diversifying rather than convergent evolution (de Guillen et al. 2015).

Structural similarity between effectors, as with the *M. oryzae* MAX effectors, is not predictive of the molecular function. For instance, whereas AVR-Pia and AVR-Pik bind heavy metal-associated (HMA) domain-containing proteins, the MAX effector AvrPiz-t targets E3 ubiquitin ligases (Cesari et al. 2013; Park et al. 2012; Park et al. 2016; Wu et al. 2015). The MAX fold most likely provides a basic structural scaffold that confers stability in the host, while providing a framework for sequence diversification and terminal extensions that result in adaptive evolution and neofunctionalization of the effectors (de Guillen et al. 2015; Yoshida et al. 2016). Beyond MAX effectors, there is emerging evidence that other families of filamentous pathogen effectors have expanded from a common ancestor by duplication and diversification (Franceschetti et al. 2017; Win et al. 2012b). Conserved folds, such as the oomycete WY-fold, also serve as a chassis to support protein structural integrity, while providing enough plasticity for the effectors to evolve unrelated activities inside host cells (Boutemy et al. 2011).

### Lesson 5. Many effectors, one target: pathogens hedging their bets

Different effectors can target the same host protein whether the effectors originate from a single pathogen, i.e. functionally redundant effectors, or from phylogenetically unrelated pathogens (Macho and Zipfel 2014; Mukhtar et al. 2011; Song et al. 2009) (Fig. 4A). In either case, the targets of these effectors can be viewed as “hubs”—pathways that pathogens need to manipulate for successful infection.

**Figure 4.**
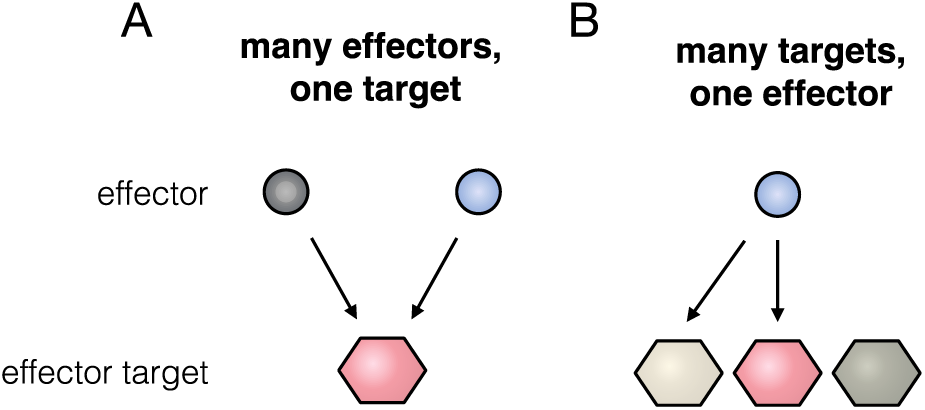
Pathogen effectors and their host targets. (A) Unrelated effector proteins can converge on the same plant target as illustrated in ‘many effectors, one target’ panel. (B) ‘Many targets, one effector’ shows that some effectors can associate with multiple members of the same protein family and discriminate between closely related host proteins.

Examples of effectors that converge on related host proteins include *M. oryzae* AVR-Pik, AVR-Pia, and AVR1-CO39. Each of these effectors bind rice proteins with HMA domains (Cesari et al. 2013; Kanzaki et al. 2012; Maqbool et al. 2015). Although the precise identity of the host targets of these effectors is not yet clear, AVR-Pik, AVR-Pia and AVR1-CO39 are thought to bind HMA-containing proteins in order to promote infection. Indeed, one such rice HMA-containing protein, Pi21, acts as a host susceptibility (*S*-) gene during *M. oryzae* infection (Fukuoka et al. 2009). Pi21 harbors an N-terminal HMA domain followed by a proline rich region and a C-terminal isoprenylation signal, and deletions within the proline rich domain confer genetically recessive resistance to *M. oryzae* (Fukuoka et al. 2009).

Why have effectors from the same species evolved to converge on the same host targets? One possibility is that effector redundancy is a product of arms race coevolution between pathogens and hosts. In biology, genetic redundancy contributes to robustness in the face of a changing environment. Similarly, in an evolutionary strategy referred to as “bet-hedging”, plant pathogen populations benefit from carrying a redundant reservoir of effectors to counteract host immunity (Win et al. 2012b). This could also explain why most effector proteins are not essential for host colonization, but rather have quantitative effects on pathogen virulence.

### Lesson 6. Many targets, one effector: the discriminating power of effectors

Pathogen effectors that associate with related members of a host protein family have often evolved the capacity to discriminate between closely related proteins (Fig. 4B). One example is the *M. oryzae* effector AVR-Pii, which binds only two members of the large Exo70 protein family of rice, OsExo70F-2 and OsExo70F-3 (Fujisaki et al. 2015). Exo70 is a subunit of the exocyst, an evolutionarily conserved vesicle tethering complex that functions in the last stage of exocytosis (Heider and Munson 2012; Zhang et al. 2010). In plants, Exo70 subunits have largely expanded, particularly in grasses, with 47 paralogs in rice (Cvrčková et al. 2012; Synek et al. 2006). Why AVR-Pii targets OsExo70F-2 and OsExo70F-3, and how these interactions affect pathogenesis, remains to be determined (Fujisaki et al. 2015).

There are additional examples of effectors that can discriminate between related proteins. PexRD54, an effector from the potato late blight pathogen *Phytophthora infestans*, discerns between members of the ATG8 host protein family, ubiquitin-like proteins that modulate selective autophagy networks (Dagdas et al. 2016). As with the Exo70 family, ATG8 proteins have dramatically expanded in plants (Kellner et al. 2017; Seo et al. 2016). Remarkably, PexRD54 binds one specific member of potato ATG8 family, ATG8CL, with a 10-fold higher affinity than another member, ATG8IL (Dagdas et al. 2016; Maqbool et al. 2016). This enables the pathogen to hijack a specific selective autophagy pathway in the host to promote colonization (Dagdas et al. 2017).

### Lesson 7. Beyond gene-for-gene interactions: NLRs working together

Disease resistance genes in plants often encode immune receptors of the NLR class (Jacob et al. 2013; Jones et al. 2016). Although the majority of these genes are identified as single loci that follow Flor’s gene-for-gene postulate (Flor 1971), the underlying genetic architecture of disease resistance is much more complex. NLR proteins are now known to function as pairs, or even to form intricate genetic networks (Eitas and Dangl 2010; Wu et al. 2017). The prevailing model is that sensor NLRs, which detect pathogen effectors, require helper NLRs to activate defense signaling (Bonardi et al. 2011; Cesari et al. 2014a; Eitas and Dangl 2010; Wu et al. 2016). The NLR proteins that detect AVR-Pik, AVR-Pia and AVR-Pii represent three classic examples of NLR pairs (Fig. 1). These three NLR pairs, i.e. Pik (Pik-1/Pik-2), Pia (RGA5/RGA4) and Pii (Pii-2/Pii-1), are genetically linked in head-to-head orientation, and in each case the sensor partner has integrated a non-canonical domain that mediates pathogen detection (Ashikawa et al. 2008; Lee et al. 2009; Okuyama et al. 2011) (Fig. 5).

**Figure 5.**
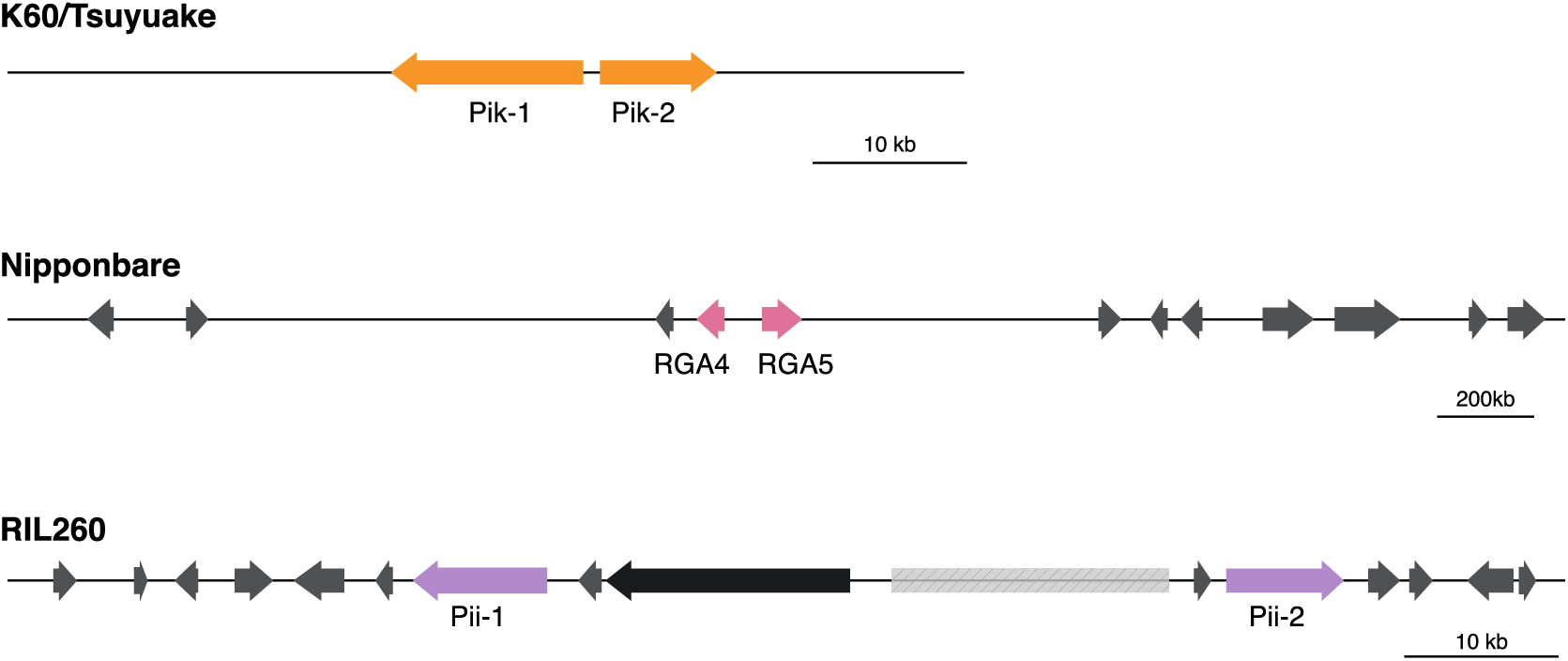
Pik, Pia and Pii NLR pairs are genetically linked. (Top) Genetic linkage of *Pik*-*1* and *Pik*-*2* genes (orange) in K60/Tsuyuake cultivar. (Middle) The *Pia* locus with *RGA4* and *RGA5* (pink) in head-to-head orientation in Nipponbare cultivar. (Bottom) *Pii*-*1* and *Pii*-*2* genes (purple) in *Pii* locus, also known as Pi5, of RIL260 rice cultivar. Other genes (grey) and transposon (black) mapped to respective loci are illustrated by arrowheads representing the direction of coding strand. A gap in the DNA sequence in RIL260 is indicated by dashed box.

It may seem surprising that all three effectors identified by Yoshida et al. (2009) turned out to have cognate NLR pairs, as they represent deviations from the classic gene-for-gene model. However, there are now many examples of paired NLRs across plant species that detect an array of pathogens (Cesari et al. 2014a; Jones and Dangl 2006). In one case, the RRS1/RPS4 NLR pair from *Arabidopsis thaliana* contributes to resistance to both bacterial and fungal pathogens (Narusaka et al. 2009). Remarkably, even classic NLRs that were long thought to function as singletons, such as the Solanaceae NLRs Prf, Bs2, Rx, and Mi-1, turned out to require NLR mates (Wu et al. 2017). In this case, these sensor NLRs belong to a large NLR network that confers immunity to oomycetes, bacteria, viruses, nematodes, and insects (Wu et al. 2017). The network has emerged over 100 Mya from a linked NLR pair that diversified into up to one-half of the NLRs of asterid plants (Wu et al. 2017).

### Lesson 8. NLR unconventional domains: there is nothing stranger in a strange NLR protein than the strange domain that’s integrated in it

NLRs are modular proteins that contain three classical domains: an N-terminal coiled-coil CC or TOLL/interleukin-1 receptor (TIR) domain, a central nucleotide binding pocket (NB), and a C-terminal leucine rich repeat (LRR) domain (Dodds and Rathjen 2010; McHale et al. 2006; Takken and Goverse 2012). Although the majority of NLR receptors share this conserved domain architecture, anywhere from 3 to 10% of plant NLRs turned out to include extraneous domains in addition to the classic NLR architecture (Kroj et al. 2016; Sarris et al. 2016). These so-called “integrated domains” are thought to have evolved from effector targets to mediate pathogen detection, either by binding effectors or by serving as substrates for the effector’s enzymatic activity (Cesari et al. 2014a; Wu et al. 2015) (Fig. 6). Remarkably, all three NLRs that detect the *M. oryzae* effectors AVR-Pik, AVR-Pia, and AVR-Pii, contain integrated domains. Indeed, the integrated heavy metal-associated (HMA) domains of both Pik-1 and RGA5 receptors directly bind AVR-Pik and AVR-Pia/AVR1-CO39, respectively (Cesari et al. 2013; Maqbool et al. 2015; Ortiz et al. 2017). It’s notable that a single type of NLR-integrated domain mediates perception of three evolutionarily unrelated effectors, suggesting that integration of extraneous domains into immune receptors may increase robustness of the immune response against promiscuous effectors.

**Figure 6.**
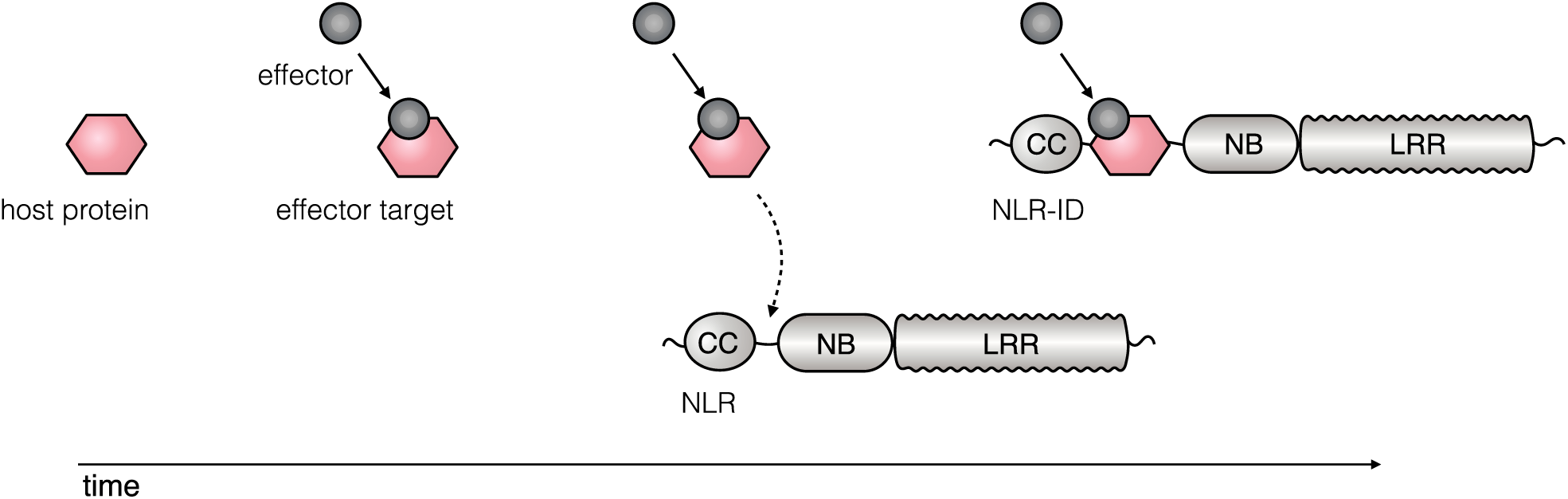
Evolution on an NLR-integrated domain. Some NLRs can carry integrated domains that are thought to be derived from the operative targets of effectors. In this model, a host protein first becomes an effector target. Subsequently, the target integrates into a host NLR where it becomes a bait for effector recognition.

The discovery that NLR proteins contain unconventional domains that have evolved by duplication of an effector target, followed by fusion into the NLR, signals the merger of two fields—NLR biology and effector biology. There are practical consequences of this conceptual development: integrated domains could serve as a source for discovery of host factors that act as hubs for pathogen effectors. Moreover, in many cases, the unintegrated paralogs of extraneous NLR-domains may encode susceptibility genes, and thus serve as platforms for crop improvement (Kroj et al. 2016; Sarris et al. 2016).

### Lesson 9. Mechanisms of immunity: NLRs dancing to different tunes

How do paired NLRs operate to activate immune signaling? A common model is that sensor NLRs hold helper NLRs in an inactive complex, and pathogen effector activity triggers the release of this negative regulation (Fig. 7A). One example of this model is the rice Pia pair. The helper NLR, RGA4, is constitutively active and triggers spontaneous cell death when expressed in the absence of its mate, RGA5 (Cesari et al. 2014b). Binding of the *M. oryzae* effectors AVR-Pia or AVR1-CO39 to RGA5 results in the release of RGA4, which leads to immune signaling and disease resistance (Cesari et al. 2013; Cesari et al. 2014b).

**Figure 7.**
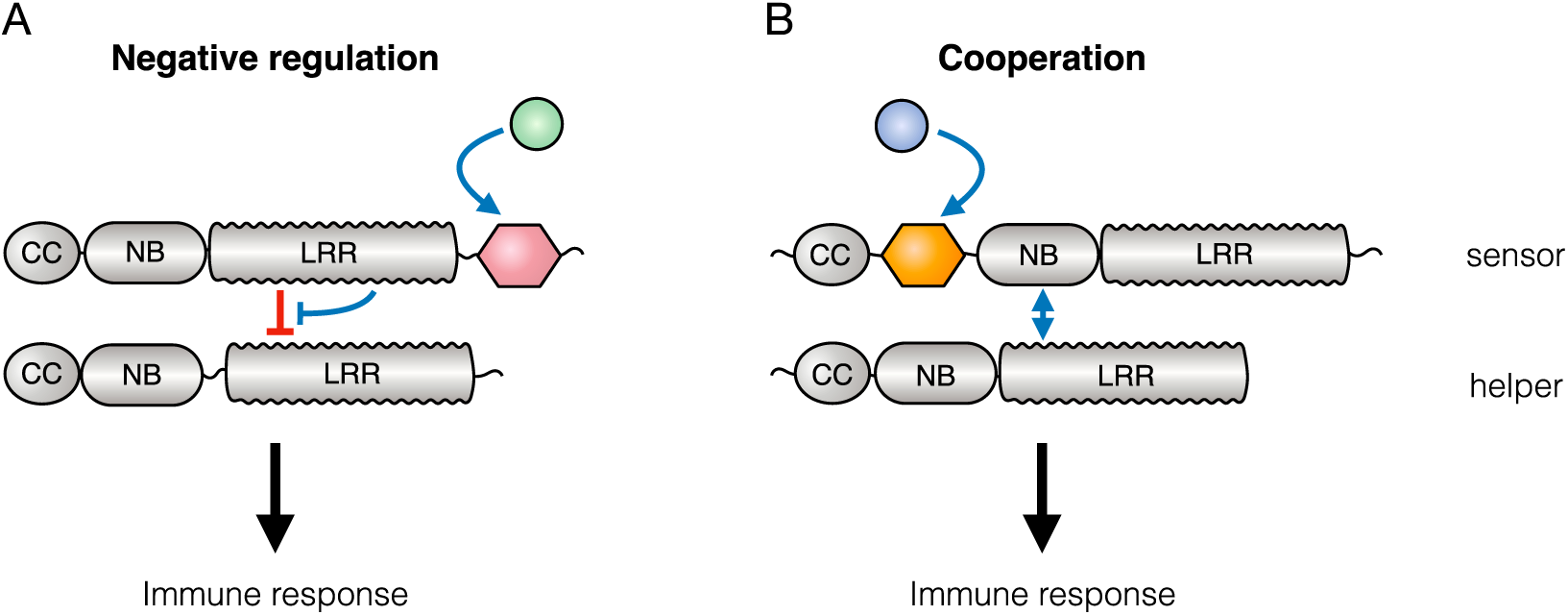
Paired NLRs can activate immunity through negative regulation or cooperation. (A) In the negative regulation model, the sensor NLR represses the autoactivity of the helper NLR in the absence of pathogen stimuli (red). Immunity is activated when the effector binds the sensor NLR, releasing the suppression of the helper NLR activity (blue). (B) In the cooperation model, sensor and helper NLRs work together to induce the immune response upon effector binding (blue). Neither the helper nor the sensor NLRs can trigger immunity alone.

Interestingly, not all NLR pairs appear to function through negative regulation. In the rice Pik pair, neither Pik-1 nor Pik-2 exhibit autoimmunity, and thus presumably operate through a different mechanism. Although the precise mechanics of the activation of the Pik pair remains to be elucidated, both NLRs are required to trigger an immune response following AVR-Pik binding to Pik-1 (Maqbool et al. 2015; Zhai et al. 2011). This suggests that the Pik pair may function through cooperation between the NLR mates (Fig. 7B).

The mechanisms by which NLR pairs function is likely to impose different constraints on the evolution of the corresponding NLR genes. NLRs that operate through negative regulation, such as Pia, are likely to function as a single unit and remain genetically linked to ensure fine-tuned co-regulation and prevent the accumulation of deleterious effects. Conversely, positively regulated NLR complexes may accommodate a certain degree of evolutionary plasticity, which in some cases could lead to expansions beyond genetically linked genes. The recently described NLR network of Wu et al. (2017) is an example of massive diversification of NLR genes that have probably originated from a single NLR pair (Wu et al. 2017). In contrast, it is difficult to envisage how negatively regulated NLR pairs could evolve into such a gene network without a significant fitness cost to the plant.

### Lesson 10. NLR activation of immunity: effector binding ain’t enough

NLRs need to recognize a pathogen effector to activate immunity and confer disease resistance. The current view is that NLR immune responses are the result of a two-part process: pathogen effector recognition and subsequent activation of immune signaling by the receptor. In many cases, recognition is achieved by direct binding of the NLR immune receptor to the effector. However, NLR binding may not always be enough to activate an effective immune reaction. One well-studied example of effector recognition is rice Pik-1, which forms a complex with the *M. oryzae* AVR-Pik effector via the integrated HMA domain (Maqbool et al. 2015). Although the HMA domain of Pik-1 can bind three different variants of AVR-Pik, only interaction with AVR-PikD results in disease resistance (Kanzaki et al. 2012; Maqbool et al. 2015). Interestingly, the binding affinity between HMA and AVR-PikD is at least 10 times stronger than the interactions with the other effector variants, indicating that the NLR-effector complex may need to reach a certain threshold of conformational stability to activate an effective immune response.

The finding that NLR binding to an AVR effector is not sufficient to activate disease resistance has implications for engineering synthetic plant immune receptors with expanded recognition spectra. Mutations across classical NLR domains, including the relatively conserved CC and NB domains, can yield sensitized “trigger-happy” receptors with markedly lower thresholds of activation (Chapman et al. 2014; Farnham and Baulcombe 2006; Giannakopoulou et al. 2015; Harris et al. 2013; Segretin et al. 2014; Stirnweis et al. 2014). However, such NLR mutants do not always translate into an effective resistance response against virulent races of pathogens (Segretin et al. 2014). Thus, we need to better understand the interplay between effector recognition and activation to achieve rational design of improved plant immune receptors (Giannakopoulou et al. 2016; Kim et al. 2016). Strategies for improving NLRs may need to combine different classes of mutations, e.g. mutations that expand effector binding spectrum, with “trigger-happy” mutations that result in a lower activation threshold (Fig. 8).

**Figure 8.**
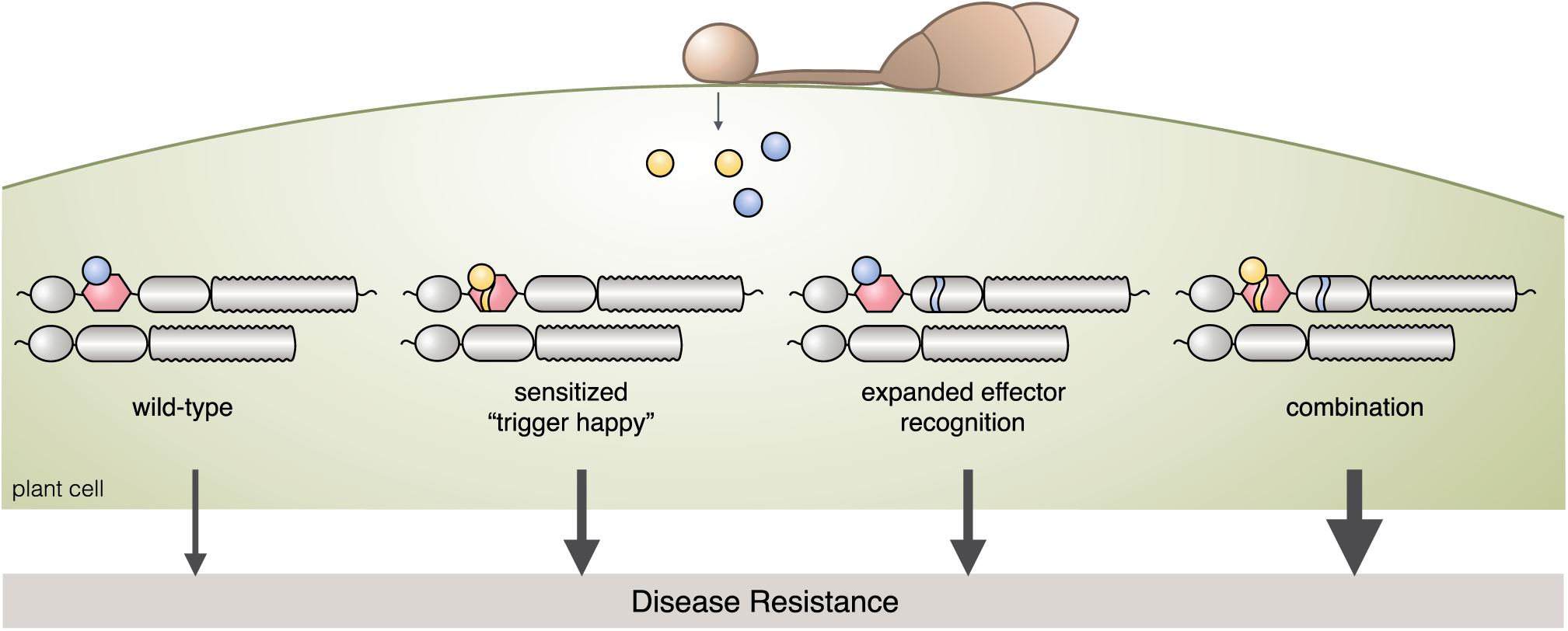
Strategies for improving NLR immune receptors. Sensitized NLR mutants are “trigger-happy,” having a lower activation threshold, whereas mutants with altered effector binding respond to a wider spectrum of effectors. We hypothesize that combining the two classes of mutants would result in more effective levels of disease resistance (illustrated by the thickness of the arrow).

### Lesson 11. Plant-pathogen arms race: NLR evolution on steroids

The coevolution of hosts and pathogens is driven by antagonistic molecular interactions in which both pathogen and plant components are under selection to adapt. As a consequence, both effector and immune receptor genes carry strong signatures of positive selection (see Lesson 3). The arms race between *M. oryzae* effector AVR-Pik and its matching rice Pik NLR demonstrates the rapid coevolution between a plant pathogen effector and an immune receptor (Fig. 9A). Patterns of *Pik*-*1* allelic diversity among rice germplasm have been interpreted as evidence that this arms race has accelerated following rice domestication (Costanzo and Jia 2010; Kanzaki et al. 2012). The most polymorphic region of *Pik*-*1* is the integrated HMA domain responsible for AVR-Pik recognition, consistent with the view that *Pik*-*1* is under strong diversifying selection imposed by the pathogen (Costanzo and Jia 2010; Zhai et al. 2014) (Fig. 9B). Interestingly, the Pik-1 helper NLR, Pik-2, exhibits lower polymorphism levels compared to Pik-1 (Kanzaki et al. 2012). The pattern of Pik-2 evolution is consistent with its putative role in immune signalling that may impose higher evolutionary constraints on the protein.

**Figure 9.**
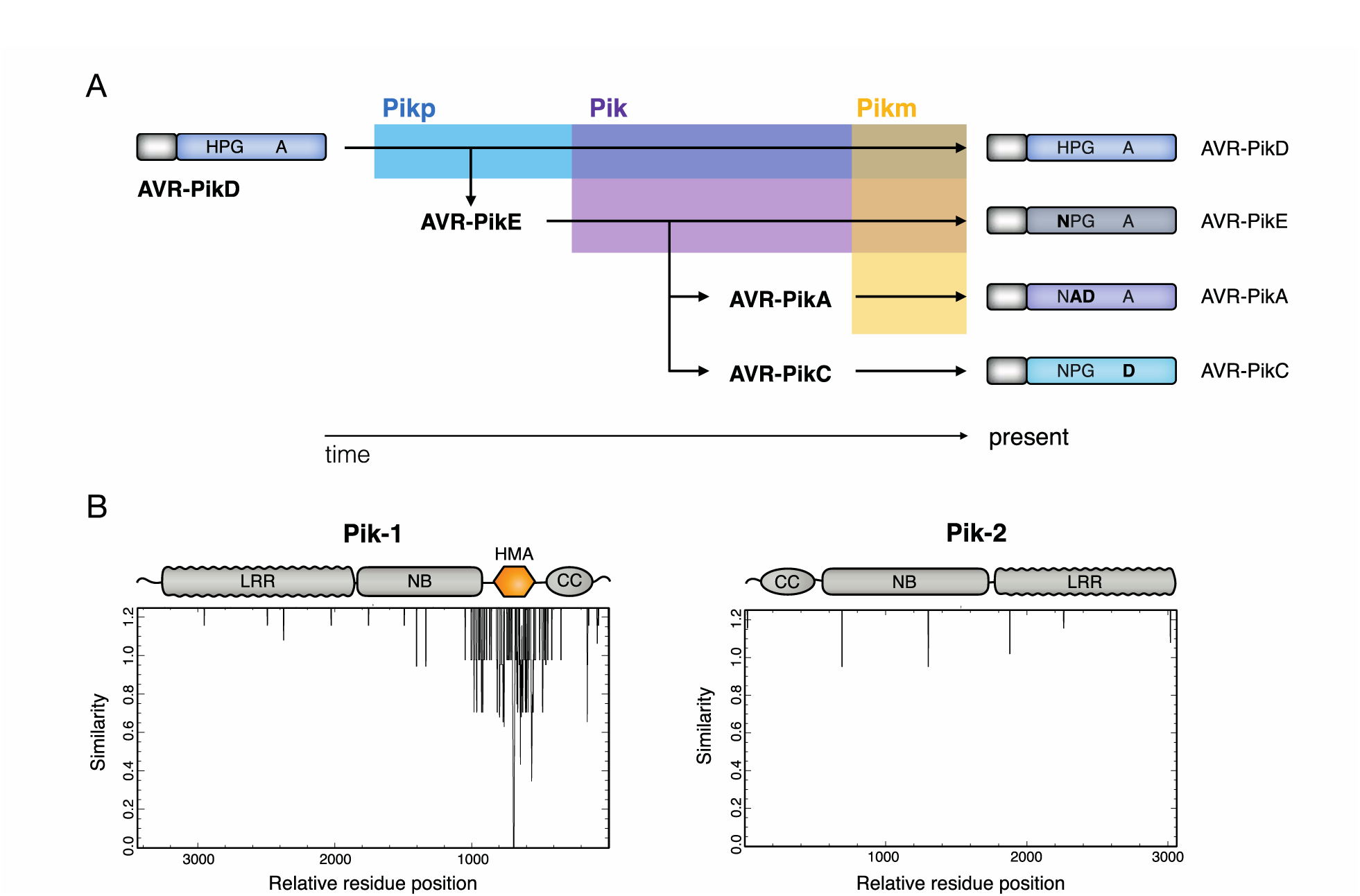
The *Magnaporthe oryzae* effector AVR-Pik has evolved through an arms race with the rice immune receptor Pik. (A) The ancestral allele of AVR-Pik effector, AVR-PikD, is recognised by the Pikp immune receptor allele (blue). AVR-PikD evaded detection by Pikp through the introduction of a non-synonymous nucleotide polymorphism, creating the AVR-PikE allele. The subsequent evolution of Pik (purple) and Pikm (yellow), which have expanded recognition spectrums, led to further non-synonymous substitutions and the deployment of AVR-PikA and AVR-PikC alleles. The nucleotide polymorphisms (bold) driven by the recognition specificity of the rice immune receptors, ultimately led to the four present-day AVR-Pik alleles. (B) Genetic diversity of *Pik*-*1* and *Pik*-*2* alleles illustrated by conservation plots of the CDS sequence alignment of 13 Pik-1 and Pik-2 alleles. The panel has been adapted from Costanzo and Jia (2010).

### Lesson 12. Into the wild: drivers of pathogen specialization in the rice terraces of Yuanyang

We understand little about the dynamics of plant and pathogen populations in the field. The Yuanyang terraces in Yunnan, China, offer an opportunity to study rice and blast fungus coevolutionary interactions in a traditional agricultural system where japonica and indica rice have been cultivated side-by-side for centuries (He et al. 2011). In Yuanyang, *M. oryzae* populations are specialized on indica and japonica rice varieties. Moreover, the japonica and indica rice isolates of *M. oryzae* differentially infected different hosts under controlled conditions, with reduced virulence of indica isolates compared with japonica isolates on japonica rice and no observed infection of japonica isolates on indica rice. The specificity of *M. oryzae* indica and japonica isolates could be explained by the complex interplay between the pathogen effector complements and variation in host immunity. Japonica-borne isolates carry larger numbers of avirulence effectors compared to the indica-borne isolates. These differences in effector repertoires correlate with considerable differences in immunity between indica and japonica rice varieties (Fig. 10). Japonica rice possess high basal immunity, and a low complement of disease resistance genes (such as NLRs), whereas indica rice has lower basal immunity and a higher complement of resistance genes. Thus, the elevated number of effectors in *M. oryzae* japonica isolates may overcome the increased basal immunity of japonica cultivars but carry a fitness cost on indica rice by triggering disease resistance. One *M. oryzae* effector, AVR-Pia, follows this pattern. AVR-Pia was found in nearly all Yuanyang japonica isolates but was absent in all indica isolates tested. In addition, AVR-Pia enhanced *M. oryzae* infection of japonica rice under controlled conditions.

**Figure 10.**
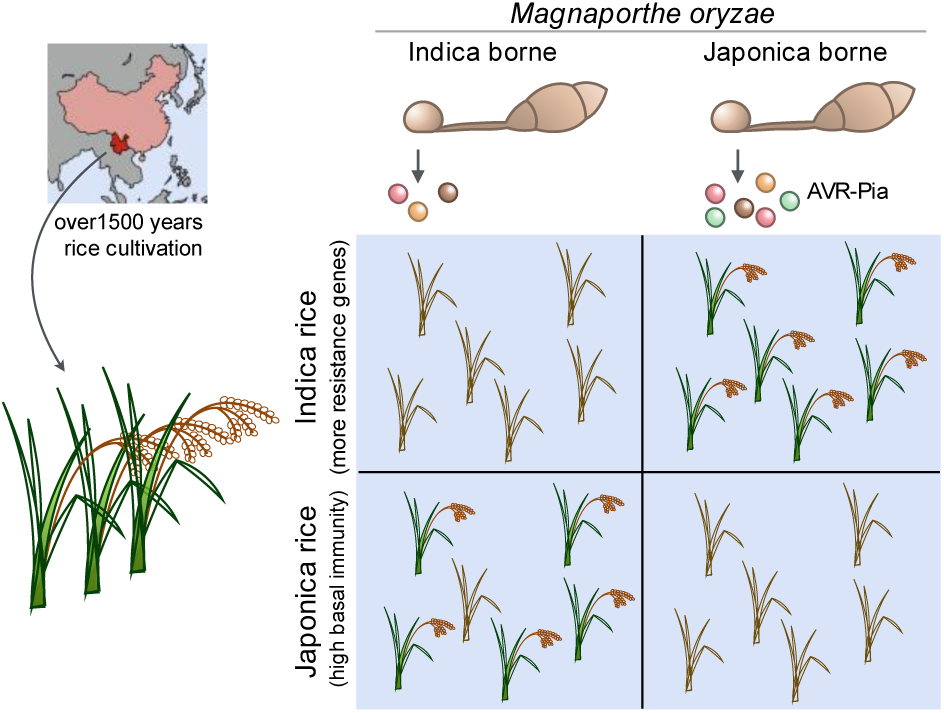
AVR-Pia into the wild. *Magnaporthe oryzae* is able to infect both indica and japónica rice. They occur side-by-side in the Yuanang terraces (map, top left), where rice has been grown for centuries. Japonica-borne isolates deploy greater numbers of effectors, allowing them to overcome the increased basal immunity of japonica rice (bottom right population panel). Indica-borne isolates deploy fewer resistance genes, allowing the fungus to evade detection by indica rice’s higher complement of resistance genes (top left population panel). Indica borne varieties were sometimes, although rarely, found on japonica rice in the field (bottom left population panel). In contrast, japonica borne varieties were largely excluded from indica rice in the field (top right population panel).

The findings from the Yuanyang terrace ecosystems have interesting implications for the management of agro-ecosystems. They demonstrate how variation in host basal- and effector-triggered immunity in a crop can lead to pathogen specialization, which in turn may limit pathogen spread on the crop and lessen destructive outbreaks. The use of mixed cropping techniques in Yuanyang contrasts sharply with the emergence of wheat blast in Brazil in the mid-1980s, which resulted from the widespread deployment of wheat varieties lacking the *RWT3* disease resistance gene and susceptible to *M. oryzae* isolates from *Lolium* spp. (Inoue et al. 2017). These *rwt3* wheat varieties served as a springboard, enabling *M. oryzae* to jump hosts to common wheat, ultimately resulting in sustained losses on this crop in South America and more recently in Asia (Inoue et al. 2017; Islam et al. 2016). Better consideration of the ecological factors associated with agricultural systems should help mitigate future outbreaks of *M. oryzae*, but also of other epidemic plant pathogens.

## CONCLUSIONS

In 2009, Hogenhout et al. (2009) reviewed the effector biology field and highlighted some of the common concepts that have stemmed from the study of plant pathogen effectors. The same year also saw Yoshida et al. (2009) describe the three AVR effectors of the rice blast fungus that inspired this article. Much has happened since. The majority of the 12 lessons we draw here describe novel concepts and insights that have emerged since Hogenhout et al. (2009) to further shape our fundamental understanding of plant-microbe interactions. Indeed, as our lessons illustrate, plant pathogen systems are great models to study evolutionary adaptations and mechanisms not just from the pathogen side but also on the plant side. Among the most dramatic conceptual developments of recent years is a better appreciation of the complexity of NLR protein structure and function. Throughout evolutionary time, NLRs have evolved to work in pairs and even networks to counteract a diversity of rapidly evolving plant pathogens (Lesson 7), and have also acquired unconventional domains to bait effectors (Lesson 8). Clearly, there are further integrated insights into plant-microbe interactions to come from the convergence of the effector biology and plant immunity fields.

We cannot underestimate the potential of translating fundamental knowledge, such as the concepts discussed in this paper, into practical applications that can solve real-world agricultural problems (Alfred et al. 2014; Kamoun 2014; Michelmore et al. 2017). There is a sense that as the science of plant-microbe interactions continues to mature and coalesce around robust principles, opportunities for practical applications are accelerating (see Supplementary Table S1 in Michelmore et al. 2017). The blast fungus is a prime example. *M. oryzae*, despite its Linnean name, is a general cereal killer that infects >50 species of grasses, including the cereal crops wheat, barley and rice (Chiapello et al. 2015; Gladieux et al. 2017; Yoshida et al. 2016). The emergence of wheat blast as a pandemic disease caused by *M. oryzae* reminds us of the importance of our science to society and the propensity of highly-adaptable pathogens to create new problems in agriculture (Inoue et al. 2017; Islam et al 2016). But there is more. The majority of the lessons drawn here are applicable to a wide-spectrum of plant-associated organisms, from symbiotic bacteria, fungi, and oomycetes to important plant pests such as insects and nematodes. Indeed, another remarkable development of the last years is the spread of the effector concept to a wide range of plant-associated organisms (Hogenhout et al. 2011; Presti et al. 2015).

To conclude, the story of these three little effectors is also a personal journey of collaboration, friendship and discovery for several of the authors of this article. It highlights the benefits of multicultural cooperation to advance science. It is gratifying that a paper published 8 years ago has served as a foundation for so much research and discovery. As our respective laboratories are nowadays fully immersed in blast fungus work, we are looking forward to many more years of exciting research.

## ACKNOWLEDGMENTS

We thank Sebastian Schornack and his own little effectors, and Robert A. Heinlein for inspiration. Work in our laboratories is funded by the Gatsby Charitable Foundation, The John Innes Foundation, Biotechnology and Biological Sciences Research Council (P012574, M02198X, M022315, P023339), European Research Council (NGRB and BLASTOFF), and Japan Society for the Promotion of Science (JSPS) Kakenhi 15H05779.

## REFERENCES

Alfred, J., Dangl, J. L., Kamoun, S., and McCouch, S. R. 2014. New Horizons for Plant Translational Research. PLOS Biol. 12:e1001880.

Allen, R.L., Bittner-Eddy, P.D., Grenville-Briggs, L.J., Meitz, J.C., Rehmany, A.P., Rose, L.E., and Beynon, J.L. 2004. Host-parasite coevolutionary conflict between Arabidopsis and downy mildew. Science 306:1957–1960.

Ashikawa, I., Hayashi, N., Yamane, H., Kanamori, H., Wu, J., Matsumoto, T., Ono, K., and Yano, M. 2008. Two adjacent nucleotide-binding site-leucine-rich repeat class genes are required to confer Pikm-specific rice blast resistance. Genetics 180:2267–2276.

Bebber, D. P., Holmes, T., and Gurr, S. J. 2014. The global spread of crop pests and pathogens. Global Ecol. Biogeogr. 23:1398–1407.

Bebber, D. P. and Gurr, S. J. 2015. Crop-destroying fungal and oomycete pathogens challenge food security. Fungal Genet. Biol. 74:62–4.

Bonardi, V., Tang, S. J., Stallmann, A., Roberts, M., Cherkis, K., and Dangl, J. L. 2011. Expanded functions for a family of plant intracellular immune receptors beyond specific recognition of pathogen effectors. Proc. Natl. Acad. Sci. USA 108:16463–16468.

Boutemy, L. S., King, S. R. F, Win, J., Hughes, R. K., Clarke, T. A., Blumenschein, R. M. A., Kamoun, S., and Banfield, M. 2011. Structures of *Phytophthora* RXLR effector proteins: a conserved but adaptable fold underpins functional diversity. J. Biol. Chem. 41:35834–42.

Cesari, S., Thilliez, G., Ribot, C., Chalvon, V., Michel, C., Jauneau, A., Rivas, S., Alaux, L., Kanzaki, H., Okuyama, Y., Morel, J.B., Fournier, E., Tharreau, D., Terauchi, R., and Kroj, T. 2013. The rice resistance protein pair RGA4/RGA5 recognizes the *Magnaporthe oryzae* effectors AVR-Pia and AVR1-CO39 by direct binding. Plant Cell 25:1463–1481.

Cesari, S., Bernoux, M., Moncuquet, P., Kroj, T., and Dodds, P.N. 2014a. A novel conserved mechanism for plant NLR protein pairs: the “integrated decoy” hypothesis. Front. Plant Sci. 5:606.

Cesari, S., Kanzaki, H., Fujiwara, T., Bernoux, M., Chalvon, V., Kawano, Y., Shimamoto, K., Dodds, P., Terauchi, R., and Kroj, T. 2014b. The NB-LRR proteins RGA4 and RGA5 interact functionally and physically to confer disease resistance. EMBO J. 33:1941–1959.

Chapman, S., Stevens, L. J., Boevink, P. C., Engelhardt, S., Alexander, C. J., Harrower, B., Champouret, N., McGeachy, K., Van Weymers, P. S. M., Chen, X., Birhc, R. J., and Hein, I. 2014. Detection of the virulent form of AVR3a from *Phytophthora infestons* following artificial evolution of potato resistance gene R3a. PLOS One 9:e110158.

Chiapello, H., Mallet, L., Guérin, c., Aguileta, G., Amselem, J., Kroj, T., Ortega-Abboud, E., Lebrun, M.H., Henrissat, B., Gendrault, A., Rodolphe, F., Tharreau, D., Fournier, E. 2015. Deciphering genome content and evolutionary relationships of isolates from the fungus *Magnaporthe oryzae* attacking different host plants. Genome Biol. Evol. 7:2896–2912.

Costanzo, S. and Jia, Y. L. 2010. Sequence variation at the rice blast resistance gene Pi-km locus: implications for the development of allele specific markers. Plant Sci. 178:523–530.

Croll, D. and McDonald, B. A. 2012. The accessory genome as a cradle for adaptive evolution in pathogens. PLOS Pathog. 8:e1002608.

Cvrčková, F., Grunt, M., Bezvoda, R., Hála, M., Kulich, I, Rawat, A., and Zársky, V. 2012. Evolution of the land plant exocyst complexes. Front. Plant Sci. 3:159.

de Guillen, K., Ortiz-Vallejo, D., Gracy, J., Fournier, E., Kroj, T., and Padilla, A. 2015. Structure analysis uncovers a highly diverse but structurally conserved effector family in phytopathogenic fungi. PLOS Pathog. 11:e1005228.

Dagdas, Y. F., Belhaj, K., Maqbool, A., Chaparro-Garcia, A., Pandey, P., Petre, B., Tabassum, N., Cruz-Mireles, N., Hughes, R. K., Sklenar, J., Win, J., Menks, F., Findlay, K., Banfield, M. J., Kamoun, S., and Bozkurt, T. O. 2016. An effector of the Irish potato famine pathogen antagonizes a host autophagy cargo receptor. eLife. 5:e10856.

Dagdas, Y., Pandey, P., Sanguankiattichai, N., Tumtas, Y., Belhaj, K., Duggan, C., Segretin, M. E., Kamoun, S., and Bozkurt, T. O. 2017. Host autophagosomes are diverted to a plant-pathogen interface. bioRxiv doi: http://dx.doi.org/10.1101/102996.

Dean, R. A., Talbot, N. J., Ebbole, D. J., Farman, M. L., Mitchell, T. K., Orbach, M. J., Thon, M., Kulkarni, R., Xu, J. R., Pan, H., Read, N. D., Lee, Y. H., Carbone, I., Brown, D., Oh, Y. Y., Donofrio, N., Jeong, J. S., Soanes, D. M., Djonovic, S., Kolomiets, E., Rehmeyer, C., Li, W., Harding, M., Kim, S., Lebrun, M. H., Bohnert, H., Coughlan, S., Butler, J., Calvo, S., Ma, L. J., Nicol, R., Purcell, S., Nusbaum, C., Galagan, J. E., and Birren, B. W. 2005. The genome sequence of the rice blast fungus *Magnaporthe grisea*. Nature 434:980–986.

Dean, R., Van Kan, J.A.L., Pretorius, Z.A., Hammond-Kosack, K.E., Di Pietro, A., Spanu, P.D., Rudd, J.J., Dickman, M., Kahmann, R., Ellis, J. and Foster, G.D. 2012. The top 10 fungal pathogens in molecular plant pathology. Mol. Plant Pathol. 13:414–430.

Dodds, P. N., Lawrence, G. J., Catanzariti, A. M., Teh, T., Wang, C. I., Ayliffe, M. A., Kobe, B., and Ellis, J. G. 2006. Direct protein interaction underlies gene-for-gene specificity and coevolution of the flax resistance genes and flax rust avirulence genes. Proc. Natl. Acad. Sci. USA. 103:8888–8893.

Dodds, P. N., and Rathjen, J. P. 2010. Plant immunity: Towards an integrated view of plant-pathogen interactions. Nat. Rev. Genet. 11:539–548.

Dong, S., Raffaele, S., and Kamoun, S. 2015. The two-speed genomes of filamentous pathogens: waltz with plants. Curr. Opin. Gen. Dev. 35:57–65.

Eitas, T.K., and Dangl, J.L. 2010. NB-LRR proteins: pairs, pieces, perception, partners, and pathways. Curr. Opin. Plant Biol. 13:472–477.

Faino, L., Seidl, M. F., Shi-Kunne, X., Pauper, M., van den Berg, G. C. M., Wittenberg, A. H. J., and Thomma, B. P. H. J. 2016. Transposons passively and actively contribute to evolution of the two-speed genome of a fungal pathogen. Genome Res. 26:1091–100.

Farhman, G., and Baulcombe, D. C. 2006. Artificial evolution extends the spectrum of viruses that are targeted by a disease-resistance gene from potato. Proc Natl Acad Sci USA 103:18828–33.

Fisher, M. C., Henk, D. A., Briggs, C. J., Brownstein, J. S., Madoff, L. C., McCraw, S. L., and Gurr, S. J. 2012. Emerging fungal threats to animal, plant and ecosystem health. Nature 484:186–194.

Flor, H. H. 1971. Current status of the gene-for-gene concept. Annual Rev. Phytopathol. 9:275–296.

Franceschetti, M., Maqbool, A., Jiménez-Dalmaroni, M. J., Pennington, H. G., Kamoun, S., and Banfield, M. J. 2017. Effectors of filamentous plant pathogens: commonalities amid diversity. Microbiol. Mol. Biol. Rev. 81: e00066–16.

Fujisaki, K., Abe, Y., Ito, A., Saitoh, H., Yoshida, K., Kanzaki, H., Kanzaki, E., Utsushi, H., Yamashita, T., Kamoun, S., and Terauchi, R. 2015. Rice Exo70 interacts with a fungal effector, AVR-Pii, and is required for AVR-Pii-triggered immunity. Plant J. 83:875–887.

Fukuoka, S., Saka, N., Koga, H., Ono, K., Shimizu, T., Ebana, K., Hayashi, N., Takahashi, A., Hirochika, H., Okuno, K., and Yano, M. 2009. Loss of function of a proline-containing protein confers durable disease resistance in rice. Science 325:998–1001.

Giannakopoulou, A., Steele, J. F. C., Segretin, M. E., Bozkurt, T. O., Zhou, J., Robatzek, S., Banfield, M. Pais, M., and Kamoun, S. 2015. Tomato I2 immune receptor can be engineered to confer partial resistance to the oomycete *Phytophthora infestans* in addition to the fungus *Fusarium oxysporum*. Mol. Plant-Microbe Interact. 28: 1316–29.

Giannakopoulou, A., Bialas, A., Kamoun, S., and Vleeshouwers, V. G. A. A. 2016. Plant immunity switched from bacteria to virus. Nat. Biotech. 34:391–392.

Gladieux, P., Condon, B., Ravel, S., Soanes, D., Nunes Maciel, J. L., Nhani Jr. A., Terauchi, R., Lebrun, M. H., Tharreau, D., Mitchell, T., Pedley, K.F., Valent, B., Talbot, N., Farman, M., and Fournier, E. 2017. Gene flow between divergent cereal- and grass-specific lineages of the rice blast fungus *Magnaporthe oryzae*. BioRxiv doi: https://doi.org/10.1101/161513.

Haas, B.J., Kamoun, S., Zody, M.C., Jiang, R.H.Y., Handsaker, R.E., Cano, L.M., Grabherr, M., Kodira, C.D., Raffaele, S., Torto-Alalibo, T., Bozkurt, T.O., Ah-Fong, A.M.V., Alvarado, L., Anderson, V.L., Armstrong, M.R., Avrova, A.O., Baxter, L., Beynon, J.L., Boevink, P.C., Bollmann, S.R., Bos, J.I.B., Bulone, V., Cai, G., Cakir, C., Carrington, J.C., Chawner, M., Conti, L., Costanzo, S., Ewan, R., Fahlgren, N., Fischbach, M.l.A., Fugelstad, J., Gilroy, E.M., Gnerre, S., Green, P.J., Grenville-Briggs, L.J., Griffith, J.M., Grünwald, N.J., Horn, K., Horner, N.R., Hu, C.H., Huitema, E., Jeong, D.H., Jones, A.M.E., Jones, J.D.G. Jones, R.W., Karlsson, E.K., Kunjeti, S.G., Lamour, K., Liu, Z., Ma, L.J., Maclean, D.J., Chibucos, M.C., McDonald, H., McWalters, J., Meijer, H.J.G., Morgan, W., Morris, P.F., Munro, C.A., O’Neill, K., Ospina-Giraldo, M.D., Pinzon, A., Pritchard, L., Ramsahoye, B., Ren, Q., Restrepo, S., Roy, S., Sadanandom, A., Savidor, A., Schornack, S., Schwartz, D.C., Schumann, U.D., Schwessinger, B., Seyer, L., Sharpe, T., Silvar, C., Song, J., Studholme, D.J., Sykes, S., Thines, M., van de Vondervoort, P.J.I., Phuntumart, V., Wawra, S., Weide, R., Win, J., Young, C., Zhou, S., Fry, W.E., Meyers, B.C., van West, P., Ristaino, J.B., Govers, F., Birch, P.R.J., Whisson, S.C., Judelson, H.S. and Nusbaum, C. 2009. Genome sequence and analysis of the Irish potato famine pathogen *Phytophthora infestans*. Nature 461:393–398.

Harris, C. J., Slootweg, E. J, Goverse, A, and Baulcombe, D. C. 2013. Stepwise artificial evolution of a plant disease resistance gene. Proc Natl Acad Sci USA 110:21189–94.

He, X., Sun, Y., Gao, D., Wei, F., Pan, L., Guo, C., mao, R., Xie, Y., Chengyun, L., and Youyong, Z. 2011. Comparison of agronomic traits between rice landraces and modern varieties at different altitudes in the paddy fields of Yuanyang terrace, Yunnan province. J. Resour. Ecol. 2:46–50.

Heider, M. R. and Munson, M. 2012. Exorcising the exocyst complex. Traffic. 13: 898–907.

Hogenhout, S. A., Van der Hoorn, R. A. L., Terauchi, R., and Kamoun, S. 2009. Emerging concepts in effector biology of plant-associated organisms. Mol. Plant-Microbe. Interact. 22:115–22

Hoegenhout, S. A., and Bos, J. I.B. 2011. Effector proteins that modulate plant-insect interactions. Curr. Opin. Plant Biol. 14:422–428.

Huang, J., Si, W., Deng, Q., Li, P., and Yang, S. 2014. Rapid evolution of avirulence genes in rice blast fungus *Magnaporthe oryzae*. BMC Genetics 15:45.

Inoue, Y., Vy, T. T. P., Yoshida, K., Asano, H., Mitsuoka, C., Asuke, S., Anh, V. L., Cumagun, C. J. R., Chuma, I., Terauchi, R., Kato, K., Mitchell, T., Valent, B., Farman, M., and Tosa, Y. 2017. Evolution of the wheat blast fungus through functional losses in a host specificity determinant. Science 357:80–83.

Islam, M. T., Croll, D., Gladieux, P., Soanes, D. M., Persoons, A., Bhattacharjee, P., Hossain, M. S., Gupta, D. R., Rahman, M. M., Mahboob, M. G., Cook, N., Salam, M. U., Surovy, M. Z., Sancho, V. B., Maciel, J. L., NhaniJunior, A., Castroagudin, V. L., Reges, J. T., Ceresini, P. C., Ravel, S., Kellner, R., Fournier, E., Tharreau, D., Lebrun, M. H., McDonald, B. A., Stitt, T., Swan, D., Talbot, N. J., Saunders, D. G., Win, J., and Kamoun, S. 2016. Emergence of wheat blast in Bangladesh was caused by a South American lineage of *Magnaporthe oryzae*. BMC Biol 14:84.

Jacob, F., Vernaldi, S., and Maekawa, T. 2013. Evolution and conservation of plant NLR functions. Front. Immunol. 4:297.

Jones, J. D., and Dangl, J. L. 2006. The plant immune system. Nature 444:323–329.

Jones, J. D., Vance, R. E., and Dangl, J.L. 2016. Intracellular innate immune surveillance devices in plants and animals. Science 354:6316.

Kamoun, S. 2014. Gained in translation. IS-MPMI Reporter 1:2.

Kanzaki, H., Yoshida, K., Saitoh, H., Fujisaki, K., Hirabuchi, A., Alaux, L., Fournier, E., Tharreau, D., and Terauchi, R. 2012. Arms race co-evolution of *Magnaporthe oryzae* AVR-Pik and rice Pik genes driven by their physical interactions. Plant J. 72:894–907.

Kellner, R., De la Concepcion, J., Maqbool, A., Kamoun, S., and Dagdas, Y. F. 2017. ATG8 expansion: a driver of selective autophagy diversification? Trends Plant Sci. 22:204–214.

Kim, S. H., Qi, D., Ashfield, T., Helm, M. and Innes, R. W. 2016. Using decoys to expand the recognition specificity of a plant disease resistance protein. Science 351:684–687.

Kroj, T., Chanclud, E., Michel-Romiti, C., Grand, X., and Morel, J. B. 2016. Integration of decoy domains derived from protein targets of pathogen effectors into plant immune receptors is widespread. New Phytol. 210:618–626

Lee, S.K., Song, M.Y., Seo, Y.S., Kim, H.K., Ko, S., Cao, P.J., Suh, J.P., Yi, G., Roh, J.H., Lee, S., An, G., Hahn, T.R., Wang, G.L., Ronald, P., and Jeon, J.S. 2009. Rice Pi5-mediated resistance to *Magnaporthe oryzae* requires the presence of two coiled-coil-nucleotide-binding-leucine-rich repeat genes. Genetics 181:1627–1638.

Liu, Z., Bos, J. I. B., Armstrong, M., Whisson, S. C., da Cunha, L., Torto-Alalibo, T., Win, J., Avrova, A. O., Wright, F., Birch, P. R., and Kamoun, S. 2005. Patterns of diversifying selection in the phytotoxin-like scr74 gene family of *Phytophthora infestans*. Mol. Biol. Evol. 22:659–672.

Liu, W., Liu, J., Triplett, L., Leach, J. E., and Wang, G.-L. 2014. Novel insights into rice innate immunity against bacterial and fungal pathogens. Annu. Rev. Phytopath. 52:213–241.

MacLean, A. M., Orlovskis, Z., Kowitwanich, K., Zdziarska, A M., Angenent, G. C., Immink, R. G. H., and Hogenhout, S. A. 2014. Phytoplasma effector SAP54 hijacks plant reproduction by degrading MADS-box proteins and promotes insect colonization in a RAD23-dependent manner. PLOS Biol. 12: e1001835.

Macho, A. P., and Zipfel, C. 2015. Targeting of plant pattern recognition receptor-triggered immunity by bacterial type-III secretion system effectors. Curr. Opin. Micro. 23:14–22.

Maekawa, T., Kufer, T. A., and Schulze-Lefert, P. 2011. NLR functions in plant and animal immune systems - so far and yet so close. Nature Immunol. 12:818–826.

Maqbool, A., Saitoh, H., Franceschetti, M., Stevenson, C., Uemura, A., Kanzaki, H., Kamoun, S., Terauchi, R., and Banfield, M. 2015. Structural basis of pathogen recognition by an integrated HMA domain in a plant NLR immune receptor. eLife 4:1–24.

Maqbool, A., Hughes, R. K., Dagdas, Y. F., Tregidgo, N., Zess, E., Belhaj, K., Round, A., Bozkurt, T. O., Kamoun, S., and Banfield, M. J. 2016. Structural basis of host autophagy-related protein 8 (ATG8) Binding by the Irish potato famine pathogen effector protein PexRD54. J. Biol. Chem. 291:20270–20282.

McHale, L., Tan, X., Koehl, P., and Michelmore, R. W. 2006. Plant NBS-LRR proteins: adaptable guards. Genome Biol. 7:212.

Michelmore, R. W., Coaker, G., Bart, R., Beattie, G. A., Bent, A., Bruce, T., Cameron, D., Dangl, J., Dinesh-Kumar, S., Edwards, R. Eves-van den Akker, S., et al. 2017. Foundational and translational research opportunities to improve plant health. Mol. Plant-Microbe Interact. 30:515–516.

Mukhtar, M.S., Carvunis, A.R., Dreze, M., Epple, P., Steinbrenner, J., Moore, J., Tasan, M., Galli, M., Hao, T., Nishimura, M.T., Pevzner, S.J., Donovan, S.E., Ghamsari, L., Santhanam, B., Romero, V., Poulin, M.M., Gebreab, F., Gutierrez, B.J., Tam, S, Monachello, D., Boxem, M., Harbort, C.J., McDonald, N., Gai, L., Chen, H., He, Y.; European Union Effectoromics Consortium, Vandenhaute, J., Roth, F.P., Hill, D.E., Ecker, J.R., Vidal, M., Beynon, J., Braun, P. and Dangl, J.L. 2011. Independently evolved virulence effectors converge onto hubs in a plant immune system network. Science 333:596–601.

Narusaka, M., Shirasu, K., Noutoshi, Y., Kubo, Y., Shiraishi, T., Iwabuchi, M., and Narusaka, Y. 2009. RRS1 and RPS4 provide a dual resistance-gene system against fungal and bacterial pathogens. Plant J. 60:218–226.

Nyarko, A., Singrarapu, K. K., Figueroa, M., Manning, V. A., Pandelova, I., Wolpert, P. J., Ciuffettti, L. M., and Barbar, E. 2014. Solution NMR structures of *Pyrenophora tritici-repentis* ToxB and its inactive homolog reveal potential determinants of toxin activity J Biol. Chem. 289:25946–25956.

Okuyama, Y., Kanzaki, H., Abe, A., Yoshida, K., Tamiru, M., Saitoh, H., Fujibe, T., Matsumura, H., Shenton, M., Galam, D.C., Undan, J., Ito, A., Sone, T., and Terauchi, R. 2011. A multifaceted genomics approach allows the isolation of the rice Pia-blast resistance gene consisting of two adjacent NBS-LRR protein genes. Plant J. 66:467–479.

Ortiz, D., de Guillen, K., Cesari, S., Chalvon, V., Gracy, J., Padilla, A., and Kroj, T. 2017. Recognition of the *Magnaporthe oryzae* effector AVR-Pia by the decoy domain of the rice NLR immune receptor RGA5. Plant Cell 29:156–168.

Ose, T., Oikawa, A., Nakamura, U., Maenaka, K., Higuchi, Y., Satoh, Y., Fujwara, S., Demura, M., Sone, T., and Kamiya, M. 2015. Solution structure of an avirulence protein, AVR-Pia, from *Magnaporthe oryzae*. J Biomol. NMR 63:229–235.

Pais, M., Win, J., Yoshida, K., Etherington, G.J., Cano, L.M., Raffaele, S., Banfield, M.J., Jones, A., Kamoun, S., and Saunders, D.G.O. 2013. From pathogen genomes to host plant processes: the power of plant parasitic oomycetes. Genome Biol. 14:211.

Park, C. H., Chen, S., Shirsekar, G., Zhou, B., Khang, C. H., Songkumarn, P., Afzal, A. J., Ning, Y., Wang, R., Bellizzi, M., Valent, B., and Wang, G. L. 2012. The *Magnaporthe oryzae* effector AvrPiz-t targets the RING E3 ubiquitin ligase APIP6 to suppress pathogen-associated molecular pattern-triggered immunity in rice. Plant Cell 24:4748–4762.

Park, C. H., Shirsekar, G., Bellizzi, M., Chen, S., Songkumarn, P., Xie, X., Shi, X., Ning, Y., Zhou, B., Suttiviriya, P., Wang, M., Umemura, K., and Wang, G.-L. 2016. The E3 ligase APIP10 connects the effector AvrPiz-t to the NLR receptor Piz-t in rice. PLOS Pathog 12:e1005528.

Pennisi, E. 2010. Armed and dangerous. Science 327:804–805.

Pesti, L. L., Lanver, D., Schweizer, G., Tanaka, S., Liang, L., Tollot, M., Zuccaro, A., Reissmann, S., Kahmann, R. 2015. Fungal effectors and plant susceptibility. Annu. Rev. Plant Biol. 66:513–545.

Raffaele, S., Farrer, R.A., Cano, L.M., Studholme, D.J., MacLean, D., Thines, M., Jiang, R.H.Y., Zody, M.C., Kunjeti, S.G., Donofrio, N.M., Meyers, B.C., Nusbaum, C., and Kamoun, S. 2010. Genome evolution following host jumps in the Irish potato famine pathogen lineage. Science 330:1540–1543.

Raffaele, S., and Kamoun, S. 2012. Genome evolution in filamentous plant pathogens: why bigger can be better. Nat. Rev. Microbiol. 10:417–430.

Ravensdale, M., Bernoux, M., Ve, T., Kobe, T., Thrall, P. H., Ellis, J. G., Dodds, P. N. 2012. Intramolecular interaction influences binding of the flax L5 and L6 resistance proteins to their AvrL567 ligands. PLOS Pathog. 8:e1003004.

Rouxel, T., and Balesdent, M. H. 2017. Life, death and rebirth of avirulence effectors in a fungal pathogen of Brassica crops, *Leptosphaeria maculans*. New Phytol. 214: 526–532.

Sarris, P. F., Duxbury, Z., Huh, S. U., Ma, Y., Segonzac, C., Sklenar, J., Derbyshire, P., Cevik, V., Rallapalli, G., Saucet, S. B., Wirthmueller, L., Menke, F. L. H., Sohn, K. H., and Jones, J. D. G. 2015. A plant immune receptor detects pathogen effectors that target WRKY transcription factors. Cell 161:1089–1100.

Sarris, P. F., Cevik, V., Dagdas, G., Jones, J. D. G., and Krasileva, K. V. 2016. Comparative analysis of plant immune receptor architectures uncovers host proteins likely targeted by pathogens. BMC Biol. 14:8.

Segretin, M. E., Pais, M., Franceschetti, M., Chaparro-Garcia, A., Bos, J. I., Banfield, M. J., and Kamoun, S. 2014. Single amino acid mutations in the potato immune receptor R3a expand response to *Phytophthora* effectors. Mol. Plant-Microbe Interact. 27:624–37.

Seo, E., Woo, J., Park, E., Bertolani, S. J., Siegel, J. B., Choi, D., and Dinesh-Kumar, S. P. 2016. Comparative analyses of ubiquitin-like ATG8 and cysteine protease ATG4 autophagy genes in the plant lineage and cross-kingdom processing of ATG8 by ATG4. Autophagy 12:2054–2068.

Sharma, R., Mishra, B., Runge, F., and Thines, M. 2014. Gene loss rather than gene gain is associated with a host jump from monocots to dicots in the smut fungus *Melanopsichium pennsylvanicum*. Genome Biol. Evol. 6:2034–2049.

Sone, T., Takeuchi, S., Miki, S., Satoh, Y., Ohtsuka, K., Abe, A., and Asano, K. 2013. Homologous recombination causes the spontaneous deletion of AVR-Pia in *Magnaporthe oryzae*. FEMS Microbiol Lett. 339:102–9.

Song, J., Win, J., Tian, M., Schornack, S., Kaschani, F., Ilyas, M., van der Hoorn, R.A.L. and Kamoun, S. 2009. Two effectors secreted by unrelated eukaryotic plant pathogens target the tomato defense protease Rcr3. Proc. Natl. Acad. Sci. USA 106:1654–1659.

Steinbrenner, A. D., Goritschnig, S., and Staskawicz, B. J. 2015. Recognition and Activation Domains Contribute to Allele-Specific Responses of an Arabidopsis NLR Receptor to an Oomycete Effector Protein. PLOS Pathog. 11:1–19.

Stirnweis, D., Milani, S. D., Jordan, T., Keller, B., and Brunner, S. 2014. Substitutions of two amino acids in the nucleotide-binding site domain of a resistance protein enhance the hypersensitive response and enlarge the PM3F resistance spectrum in wheat. Mol Plant-Microbe Interact. 27: 265–76.

Synek, L., Schlager, N., Eliáš, M., Quentin, M., Hauser, M. T., and Žárský V. 2006. AtEXO70A1, a member of a family of putative exocyst subunits specifically expanded in land plants, is important for polar growth and plant development. Plant J. 48:54–72.

Takken, F. L. and Goverse, A. 2012. How to build a pathogen detector: structural basis of NB-LRR function. Curr. Opin. Plant Biol. 15:375–384.

Talbot, N. J. 2003. On the trail of a cereal killer: Exploring the biology of *Magnaporthe grisea*. Annu. Rev. Microbiol. 57:177–202.

Terauchi, R. and Yoshida, K. 2010. Towards population genomics of effector-effector target interactions. New Phytol. 187:929–39.

Van Der Hoorn, R. A. and Kamoun, S. 2008. From guard to decoy: a new model for perception of plant pathogen effectors. Plant Cell 20:2009–2017.

Win, J., Morgan, W., Bos, J., Krasileva, K. V., Cano, L. M., Chaparro-Garcia, A., Ammar, R., Staskawicz, B. J., and Kamoun, S. 2007. Adaptive evolution has targeted the C-terminal domain of the RXLR effectors of plant pathogenic oomycetes. Plant Cell, 19:2349–2369.

Win, J., Chaparro-Garcia, a, Belhaj, K., Saunders, D. G. O., Yoshida, K., Dong, S., Schornack, S., Zipfel, C., Robatzek, S., Hogenhout, S. a, and Kamoun, S. 2012a. Effector biology of plant-associated organisms: concepts and perspectives. Cold Spring Harb. Symp. Quant. Biol. 77:235–47.

Win, J., Krasileva, K. V., Kamoun, S., Shirasu, K., Staskawicz, B. J., and Banfield, M. J. 2012b. Sequence Divergent RXLR effectors share a structural fold conserved across plant pathogenic oomycete species. PLOS Pathog. 8:e1002400.

Wu, C.-H., Krasileva, K. V., Banfield, M. J., Terauchi, R., and Kamoun, S. 2015. The “sensor domains” of plant NLR proteins: more than decoys?. Front. Plant Sci. 6:5–7.

Wu, C.-H., Belhaj, K., Bozkurt, T.O., Birk, M.S., and Kamoun, S. 2016. Helper NLR proteins NRC2a/b and NRC3 but not NRC1 are required for Pto-mediated cell death and resistance in *Nicotiana benthamiana*. New Phytol. 209:1344–1352.

Wu, C.-H., Abd-El-Haliem, A., Bozkurt, T. O., Belhaj, K., Terauchi, R., Vossen, J. H., and Kamoun, S. 2017. NLR network mediates immunity to diverse plant pathogens. Proc. Natl. Acad. Sci. USA 114:8113–8118.

Xue, M., Yang, J., Li, Z., Hu, S., Yao, N., Dean, R. A., Zhou, W., Shen, M., Zhang, H., Li, C., Liu, L., Cao, L., Xu, X., Xing, Y., Hsiang, T., Zhang, Z., Xu, J.-R., and Peng, Y.-L. 2012. Comparative analysis of the genomes of two field isolates of the rice blast fungus *Magnaporthe oryzae*. PLOS Genetics 8:e1002869.

Yasuda, N., Noguchi, M. T., and Fujita, Y. 2010. Partial mapping of avirulence genes *AVR-Pii* and *AVR*-*Pia* in the rice blast fungus *Magnaporthe oryzae*. Can. J. Plant Pathol. 28:494–498.

Yoshida, K., Saitoh, H., Fujisawa, S., Kanzaki, H., Matsumura, H., Yoshida, K., Tosa, Y., Chuma, I., Takano, Y., Win., J., Kamoun, S., and Terauchi, R. 2009. Association genetics reveals three novel avirulence genes from the rice blast fungal pathogen *Magnaporthe oryzae*. Plant Cell 21:1573–1591.

Yoshida, K., Saunders, D. G. O., Mitsuoka, C., Natsume, S., Kosugi, S., Saitoh, H., Inoue, Y., Chuma, I., Tosa, Y., Cano, L. M., Kamoun, S., and Terauchi, R. 2016. Host specialization of the blast fungus *Magnaporthe oryzae* is associated with dynamic gain and loss of genes linked with transposable elements. BMC Genomics 17:370.

Zhang, Y., Liu, C.-M., Emons, A.-M. C., and Ketelaar, T. 2010. The plant exocyst. J. Integr. Plant Biol. 52:138–146.

Zhang, Z. M., Zhang, X., Zhou, Z. R., Hu, H. Y., Liu, M., Zhou, B., and Zhou, J. 2013. Solution structure of the *Magnaporthe oryzae* avirulence protein AvrPiz-t. J Biomol. NMR 55:219–223.

Zhai, C., Lin, F., Dong, Z., He, X., Yuan, B., Zeng, X., Wang, L., and Pan, Q. 2011. The isolation and characterization of Pik, a rice blast resistance gene which emerged after rice domestication. New Phytol. 189:321–334.

Zhai, C., Zhang, Y., Yao, N., Lin, F., Liu, Z., Dong, Z., Wang, L., and Pan, Q. 2014. Function and interaction of the coupled genes responsible for Pik-h encoded rice blast resistance. PLOS One 9:e98067.

